# A phase decoder for reconstructing missing muscle activity

**DOI:** 10.64898/2026.10.28.762227

**Authors:** Serhii Bahdasariants, Sergiy Yakovenko

**Affiliations:** Department of Mechanical Engineering, Carnegie Mellon University, Pittsburgh, PA, USA; Department of Human Performance, West Virginia University, Morgantown, WV, USA; Rockefeller Neuroscience Institute, West Virginia University, Morgantown, WV, USA; Department of Neuroscience, West Virginia University, Morgantown, WV, USA; Department of Mechanical and Aerospace Engineering, Benjamin M. Statler College of Engineering and Mineral Resources, West Virginia University, Morgantown, WV, USA; Department of Chemical and Biomedical Engineering, Benjamin M. Statler College of Engineering and Mineral Resources, West Virginia University, Morgantown, WV, USA

## Abstract

Upper-limb assistive devices are commonly controlled with electromyographic (EMG) signals, but nerve injury, muscle loss, or practical limits on electrode placement can leave only a few usable signals, which constrains the decoding of intended movements. Here, we introduce a decoder that reconstructs missing muscle activations from sparse EMGs using a two-stage real-time approach: (1) estimating ongoing movement phase from available signals, and (2) retrieving corresponding muscle activations from a library of pre-recorded muscle activity templates. In proof-of-concept tests, *neurologically intact* participants (N=4) performed object-transfer tasks under simulated denervation conditions. The decoder accurately estimated movement phase (median root-mean-square error [RMSE] <0.17 of movement duration) and reconstructed missing muscle activations (RMSE 0.08-0.34; correlation up to 0.97) within ∼2 ms, in a previously unseen participant. The results indicate that template-based reconstruction could leverage sparse EMG for low-latency control of assistive technology.

## Introduction

Upper-limb assistive technologies, such as functional electrical stimulation (FES) systems and robotic orthoses, are commonly controlled by electromyographic (EMG) signals from muscles that remain under voluntary control. Peripheral nerve injury, muscle loss, and practical constraints on electrode placement can leave only a few available signals. Traumatic brachial plexus injury (BPI) is a prominent example. Although occurring in approximately 1% of major trauma cases [1], it often causes profound loss of sensorimotor function in the affected upper limb, chronic neuropathic pain, long-term disability, and psychological distress [2,3]. The pattern of paralysis depends on the location and extent of nerve-root damage, with upper-plexus lesions (C5–C7) commonly affecting shoulder abduction, external rotation, and elbow flexion, and lower-trunk lesions predominantly impairing hand and wrist function [4–8].

Reconstructive microsurgery—nerve grafts, transfers, neurolysis, or free-muscle transfer—can restore limited proximal movement [9]. Nonetheless, results remain variable; even after optimal repair, many patients regain only partial strength and require prolonged rehabilitation [10,11]. Assistive technologies can augment residual capacity, but their effectiveness is constrained by the sparsity of reliable EMG signals in severe cases [12–14].

Whatever its cause, this sparsity restricts the information available to algorithms that decode motor intent and reduces the reliability of the control signals they provide to assistive technologies. Conventional decoding algorithms typically assume access to a sufficient number of informative EMG signals, so their performance deteriorates significantly when information content is reduced [15,16]. Addressing this limitation, therefore, requires decoding methods capable of inferring motor intent—specifically, predicting missing muscle activations—from a limited number of EMG signals. Several existing methods accurately predict missing activations but are not yet suitable for real-time decoding (e.g., [17–19]).

Here, we propose a decoder that predicts real-time activity in select muscles even when only a limited number of EMG signals is available. We treat the recorded EMG as partial observations of a state vector that describes the evolution of muscle activity in time. In simple movement sequences—e.g., reach toward an object, grasp it, transport it over an obstacle, and return arm and hand to initial pose—the evolution of this state vector is expected to be stereotyped (e.g., see [20]). The reaching movement on its own is a stereotyped sequence of neural actions initiating proximal limb muscles that progress to execute limb transport and contact preparation [21–23].

Therefore, the underlying muscle activity corresponding to required musculoskeletal actions appears as a stereotypical template [24–26]. As the sequence unfolds, muscle activity follows the readout of a template (e.g., see [21,27]; also see [28] for related observations). The *phase* thus corresponds to a specific state vector in a high-dimensional command space. This one-dimensional representation of limb control is similar to the readout of internal state in the central pattern generator [29]. Other convergent descending and feedback pathways may also regulate the temporal progression of the current phase and output magnitude; however, a simple representation that detects movement phase using a task-specific template can provide an uncompressed set of complex muscle activations required to generate the desired action.

In this computational proof-of-concept study, we test task-specific EMG-envelope reconstruction under controlled replay conditions. A small group of able-bodied participants performed three object transfer tasks designed to assess (1) gross finger flexion/extension, (2) precision grip, and (3) wrist pronation/supination—movements frequently prioritized by individuals with impaired arm and hand function [30]. Data from three participants were used to construct muscle activity templates for each task. We selectively masked proximal or distal arm EMG channels to emulate scenarios of missing muscle activity, tested whether the decoder could reconstruct the withheld signals from those three participants in simulated real time, and then assessed generalizability on the fourth participant whose data were not used to build the templates. By demonstrating “good-enough” real-time reconstruction, we provide evidence that our approach can overcome the sparse input bottleneck and enable intuitive control of assistive technology for individuals who lack reliable signals from every relevant muscle—whether due to paralysis, muscle loss, or practical electrode placement constraints.

## Materials and Methods

### Participants

A small group of healthy adults (N = 4; 3 males, 1 female; age = 31.8 ± 10.8 years [mean ± s.d.]; body mass = 69.3 ± 2.9 kg; height = 1.77 ± 0.04 m) participated in the study. All participants were right-handed and reported no history of musculoskeletal or neurological disorders affecting their upper limbs. Three participants (Subjects 1-3) were dedicated to establishing the muscle activity templates used by a movement sequence phase decoder, while the fourth participant (Subject 4) served as a test subject. Written informed consent was obtained from each participant prior to the study, and all experimental protocols were carried out in accordance with the guidelines and regulations approved by the West Virginia University Institutional Review Board (protocol 2210669796).

### Experimental data

The Delsys Trigno® Wireless Biofeedback System was used to measure surface EMG, linear accelerations, and object interaction forces. Sensors were affixed to 15 muscles: anterior deltoid (DeltA), posterior deltoid (DeltP), middle deltoid (DeltM), short head of biceps brachii (BicSh), long head of biceps brachii (BicLo), lateral head of triceps brachii (TriLat), long head of triceps brachii (TriLo), brachioradialis (BRR), flexor carpi radialis (FCR), flexor digitorum superficialis (FDS), flexor carpi ulnaris (FCU), extensor carpi radialis (ECR), extensor digitorum communis (EDC), extensor carpi ulnaris (ECU), and abductor pollicis brevis (APB). Prior to electrode placement, participants’ skin was cleaned with isopropyl alcohol swabs, and any excess moisture was allowed to evaporate for 30 seconds. Sensors were placed using standardized anatomical landmarks based on SENIAM guidelines [31], initially positioned by palpation and subsequently fine-tuned by visually inspecting the EMG signals during a series of controlled single joint movements (e.g., shoulder abduction/adduction, elbow flexion/extension). A 16th Delsys Trigno sensor was dedicated to recording data from a force-sensitive resistor (FSR) attached to either of the experimental objects using adhesive tape. EMG and FSR data were sampled at 1926 Hz, while accelerometer data were sampled at 148 Hz; all were transmitted wirelessly in packets of 13.5 ms to MATLAB (R2024b; The MathWorks, Inc., Natick, MA, USA).

### Experimental setup

Participants were seated in an adjustable-height ergonomic chair facing a table, with the chair height set to level each participant’s elbow with the table surface (Fig 1b). A 15 cm × 15 cm square of adhesive tape marked the designated placement zone for the right palm on the table (not shown in Fig 1b for compactness). At the start and end of each repetition, participants rested their forearm comfortably on the table with the elbow flexed at approximately 90°, the right palm within the square, and the shoulders and back in a relaxed upright posture (leftmost insert in Fig 1b). A custom wooden barrier measuring 1 cm × 15 cm × 35.5 cm (width × height × depth) was placed on the table surface roughly 40 cm in front of each participant’s midline, functioning as an obstacle over which objects had to be transferred. Two custom 3D-printed platforms (different for each of the three tasks described below) used for object placement were attached to the table symmetrically 20 cm to the right and left of the barrier (not shown in Fig 1b for compactness). Three objects—a 3D-printed cube (6.5 cm × 6.5 cm × 6.5 cm) weighted by fishing weights to achieve a total mass of 460 g, a cylindrical weight measuring 3.5 cm in height by 2.5 cm in diameter and weighing 100 g, and a 3D-printed plate (5 cm × 1.5 cm × 9 cm) with a U-shaped cut for comfortable pinching, weighted to 125 g—were selected to assess gross finger flexion / extension, thumb opposition, and wrist pronation / supination (see Fig 1a).

**Fig 1.**
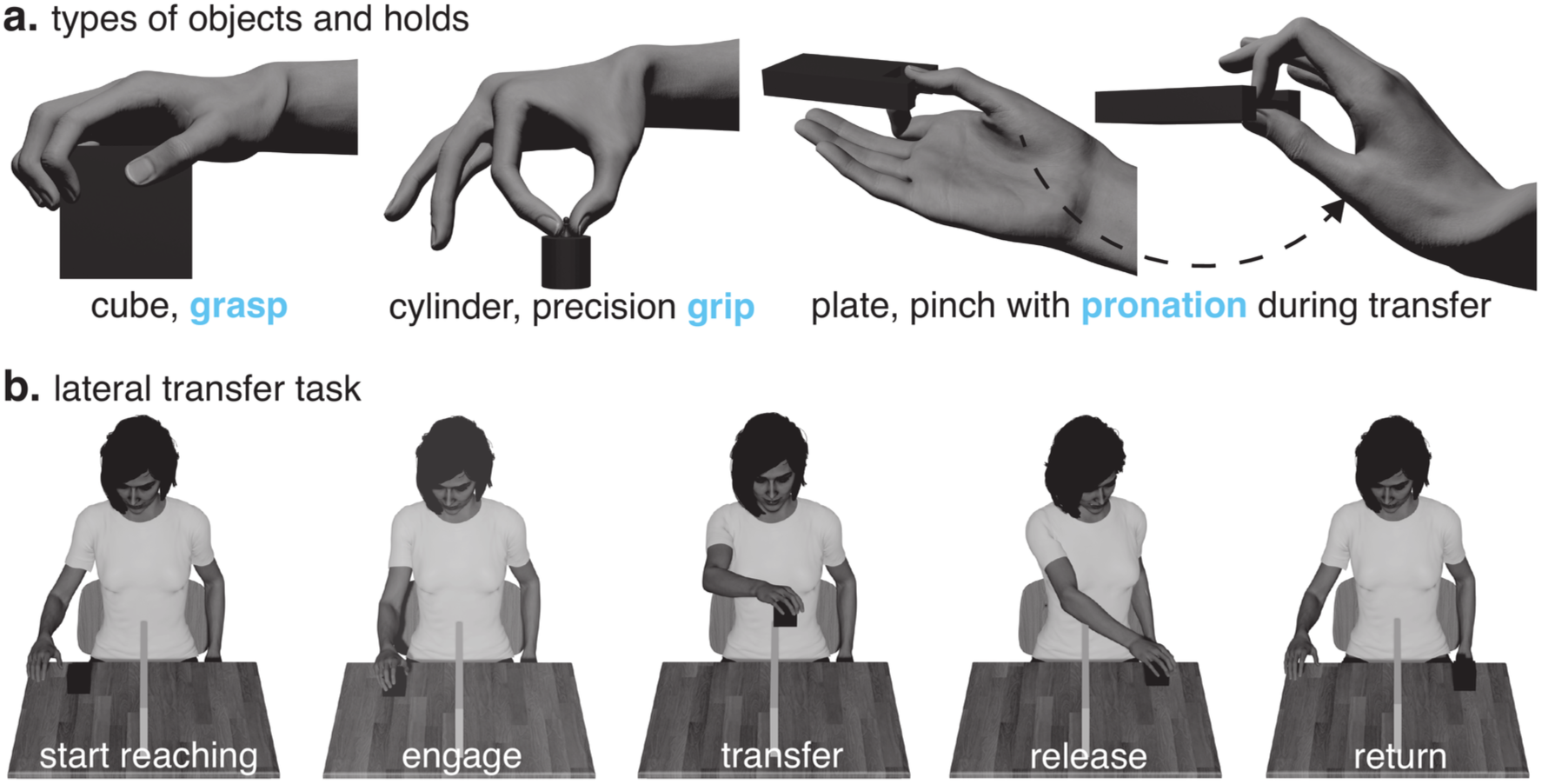
Object manipulation tasks. (**a**) The three objects used in the study—a cube (left), a cylindrical weight (middle), and a plate with a U-shaped cut (right)—are shown with hand configurations illustrating grasp (gross finger flexion-extension), grip (thumb opposition), and pinch with pronation (wrist rotation), respectively; (**b**) Representative frames of the complete movement sequence, including (from left to right) the starting posture (“start reaching”), initial object engagement (“engage”), transfer of the object (“transfer”), its release (“release”), and return of the hand to the initial resting position (“return”).

### Functional tasks

Participants performed three lateral object transfer tasks designed to assess gross finger flexion / extension (hereafter referred to as the “grasp” task), precision grip (“grip” task), and wrist pronation / supination (“pronate” task), each repeated 15 (first trial) and 30 times (second trial). To evaluate gross finger flexion-extension, participants transferred the cube (Fig 1a, leftmost panel) from one platform, over the barrier, to the opposing platform in a manner inspired by the Box and Block Test [32]. To emphasize thumb opposition, a task adapted from the Nine-Hole Peg Test [33] required pinching and placing the cylindrical weight (Fig 1a, middle panel) from one platform to the other. For wrist pronation-supination, a plate-transfer task based on a component of the Jebsen Hand Function Test [34] prompted participants to flip the plate (pronate their wrist) as they moved it across the barrier (Fig 1a, rightmost panel). In each task, participants reached for the object, grasped it, transferred it over the barrier, released it, and returned their hand to the starting position (see Fig 1b). After every repetition, the experimenter returned the object to its initial location for the subsequent repetition. A metronome set at 55 beats per minute guided the pacing of the movements; participants were instructed to synchronize their “Reach-Transfer-Return” movement sequence with each beat as closely as possible. Before data collection, three practice repetitions of each task were completed to familiarize participants with the procedure.

### Data exclusion

Post-collection, the number of repetitions per trial was standardized to 10 and 25 for the original 15- and 30-repetition trials, respectively, by excluding five repetitions from each. Exclusions were based on visually identified events characterized by: (1) aberrant changes in EMG signals indicative of artifacts, and (2) kinematic evidence of contact with the experimental barrier or interaction with the setup. When more than five such events were present, the five most prominent instances were excluded. If fewer than five were observed, the remaining exclusions proceeded sequentially from the beginning of the trial (i.e., repetition 1, followed by 2, and so on) until a total of five repetitions were removed. In Subject 4, the TriLo recordings in all three tasks and the FCR recording in the grip task contained no activity above the noise threshold (see Processing electromyography), so these two channels were excluded from the reconstruction of muscle activity; Fig 7 and 8 therefore show seven distal and six proximal muscles.

### Processing electromyography

Recorded EMG signals were processed in seven steps (Fig 2a). First, constant DC offsets were removed from task-related EMG signals by subtracting the mean signal amplitude recorded during a 30-second resting baseline period for each muscle channel (Step 1). The resulting demeaned signals were full-wave rectified (Step 2) and then smoothed using a 150 ms causal moving average filter to create a signal envelope (Step 3); window sizes of comparable duration (e.g., 200 ms) have been previously used to balance smoothness and temporal resolution of the signal (e.g., see [35]).

**Fig 2.**
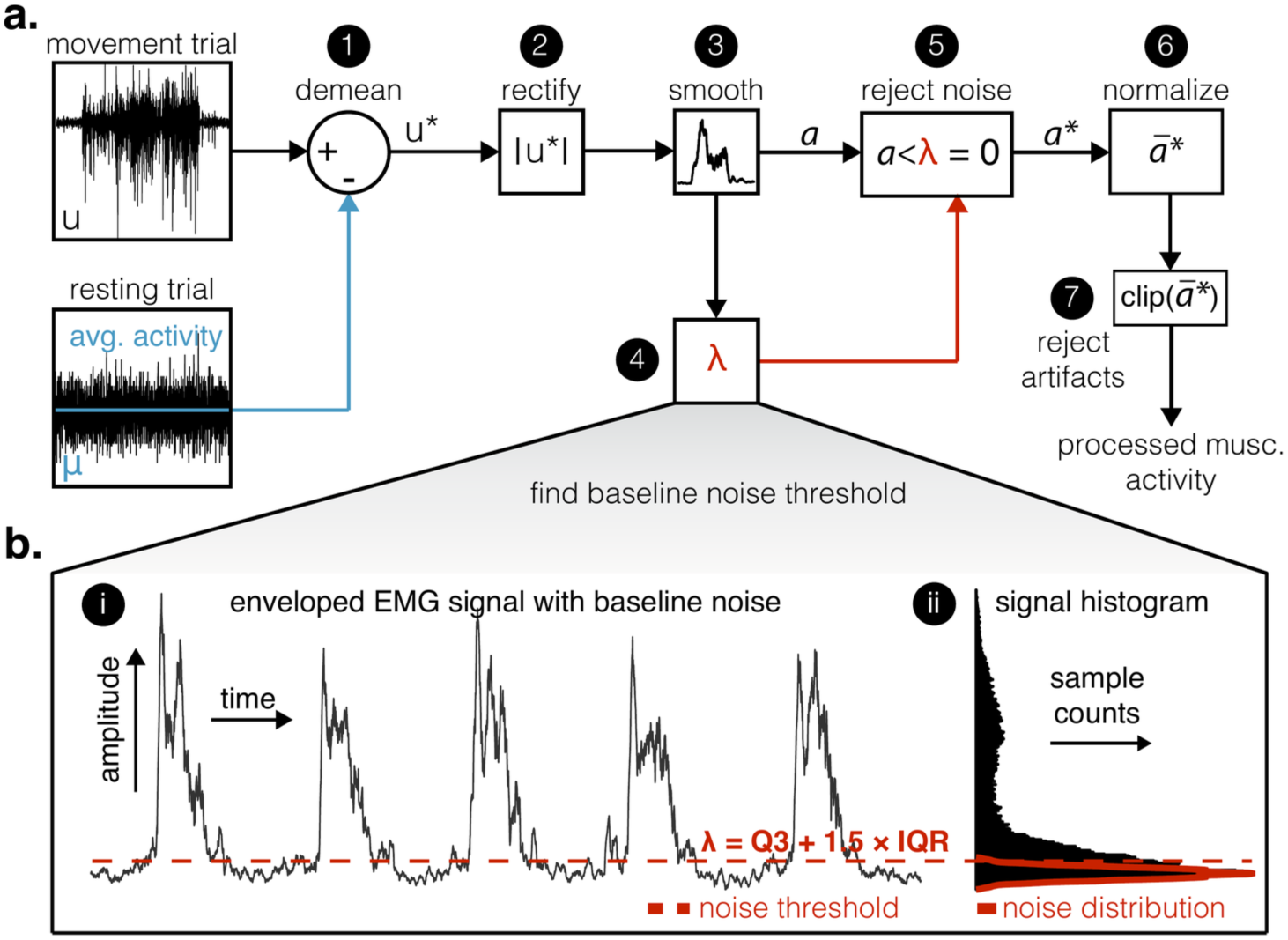
EMG processing pipeline. (**a**) Schematic of the full EMG processing pipeline. The raw EMG (*u*) from a movement trial is first demeaned (Step 1) by subtracting the average baseline activity *μ* obtained from a resting trial. The resulting signal (*u*^∗^) is then rectified (Step 2) and passed through a smoothing filter (Step 3). Next, a noise threshold *λ* is determined (Step 4) using a histogram-based approach (see panel b) and applied to remove low-amplitude components (Step 5). The resulting muscle activity *a* is normalized (Step 6) and subjected to artifact rejection (Step 7). (**b**) Magnified view illustrating how *λ* is derived. In (i), the enveloped EMG signal (black trace) still contains baseline noise (values below red dashed line). To isolate this noise, the predominant amplitude peak in the histogram (panel ii) is identified as the baseline noise level, and all amplitude values below this peak are classified as noise. Reflecting these noise values about the peak, a symmetrical estimate of the noise distribution is obtained (solid red curve), enabling calculation of the threshold (dashed red line) *λ* = Q_3_ + 1.5 · IQR (the Tukey criterion; Q_3_—third quartile; IQR — interquartile range).

To mitigate residual baseline noise, a thresholding procedure was implemented (Fig 2b, panel i). A histogram of amplitudes from the smoothed signal (output of Step 3) was generated for each muscle. The histogram mode (peak) was identified, assumed to represent the most frequent amplitude corresponding to baseline noise (Fig 2b, panel ii). The noise distribution was estimated by mirroring the sub-modal portion of the histogram across the peak value, assuming approximate symmetry (Fig 2b, panel ii, solid red curve). A muscle-specific noise rejection threshold (*λ*; Fig 2b, panel ii, dashed red line) was calculated using Tukey’s criterion: *λ* = Q_3_ + 1.5 · IQR, where Q_3_ is the third quartile and IQR is the interquartile range of this estimated noise distribution (Step 4). All smoothed EMG values falling below this threshold were set to zero (Step 5).

Finally, the processed signals underwent normalization and artifact rejection. Each denoised signal was normalized by dividing its amplitude by a subject- and muscle-specific scaling factor (Step 6) defined as the mean plus four standard deviations of the processed EMG amplitudes (output of Step 5) observed during a 30-repetition trial of each functional task. Following normalization, high-amplitude transient artifacts, potentially caused by mechanical impacts and identified as data points exceeding four standard deviations from the mean of the normalized signal, were removed (Step 7). These artifactual points were replaced using linear interpolation between the nearest valid data points. The resulting signals constituted the processed muscle activity used for generating time-normalized templates. For simulated online testing, Step 7 was excluded as real-time processing does not allow access to future data points necessary for interpolation.

### Chunking movement sequences

For each repetition of each functional task, muscle activity was computationally segmented into distinct motor chunks based on kinematic and force events. The overall movement sequence (Fig 1b) comprised a ‘Reach’ chunk, defined from movement initiation to initial object contact (first and second inserts in Fig 1b); followed by a ‘Transfer’ chunk, encompassing the period from object contact through its transfer to release (second, third, and fourth inserts in Fig 1b); and finally, a ‘Return’ chunk, covering the movement from object release back to the initial posture (fourth and fifth inserts in Fig 1b). The events marking the transitions between motor chunks were identified using data from an accelerometer and the FSR. Movement initiation (start of the ‘Reach’ chunk) was detected by the sharp increase in acceleration recorded by the sensor located over the FDS muscle. Object engagement (end of ‘Reach’ / start of ‘Transfer’ chunk) was identified by the rise in the FSR signal upon contact. Object release (end of ‘Transfer’ / start of ‘Return’ chunk) was marked by the FSR signal returning to its baseline level. Finally, the return to the initial posture (end of the ‘Return’ chunk) was determined when the acceleration of the sensor located over the FDS returned to its baseline level. Events derived from the accelerometer signal, which was pre-processed using a 2nd order low-pass Butterworth filter (10 Hz cutoff), were identified via visual inspection. FSR signal onsets and offsets were automatically detected using custom amplitude thresholding.

### Temporal normalization

Although task timing was guided by a metronome, inherent inter-subject variability was expected to lead to differences in the absolute duration of the Reach, Transfer, and Return motor chunks. To enable meaningful comparison and averaging of muscle activity templates across participants, the processed muscle activity data were time-normalized within each identified motor chunk. This involved dynamically rescaling each motor chunk (Reach, Transfer, Return) to occupy a fixed percentage of the normalized movement sequence cycle. This percentage was determined by the average relative duration of that specific motor chunk, calculated across all participants’ trials. This procedure ensured that functionally equivalent periods of muscle activation (e.g., the Reach chunk) were aligned across participants, preventing temporal misalignment in the averaged muscle activity templates. Finally, the combined length of these resampled chunks, which had a median of 6,735 samples, was subsampled to 1,000 samples. Templates were constructed from the 25-repetition trials of Subjects 1-3; the 10-repetition trials of these subjects and the 25-repetition trial of Subject 4 were reserved for testing.

### Decoding movement sequence phase and corresponding muscle activity

In an offline procedure simulating real-time processing, EMG signals were replayed in 14 ms packets, approximating the 13.5 ms packet duration of the Delsys Trigno® Wireless Biofeedback System. The original EMG data (sampled at 1926 Hz) were first processed (see Processing Electromyography) and then subsampled sixfold (∼321 Hz). This ratio was obtained from the process of subsampling the movement sequences to 1000 samples for template construction (factor ∼6.7, floored to 6). Preliminary tests (data unreported) indicated that the resulting ∼3 ms sampling interval reliably captured muscle activation envelopes. (Although these parameters sufficed for the current decoding task, we recommend further sensitivity analyses to optimize the subsampling ratio and packet duration for minimal latency and high accuracy in future real-time implementations).

Because each subsampled packet contained only four to five samples (14 ms × 1926 Hz / 6), the preceding 36 ms of muscle activity data were appended to form a 50 ms decoding window (16-17 total samples). At each decoding step, this window was slid across a 150 ms segment of the reference template, centered around the previously decoded sample index *k_t_*_–1_ (initialized at the first sample). To decode phase, the muscle activity window was shifted in integer steps *j* over the template segment; at each shift, a dot product quantified the similarity between the two:

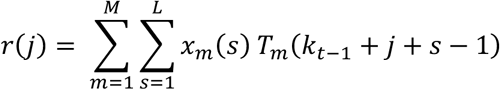

where *x_m_*(*s*) is the muscle activity from channel *m* at sample *s*, *T_m_* is the corresponding template muscle activity from the same channel, and *k_t_*_–1_ is the previously decoded template index. The subtraction of one accounts for index alignment. The shift *j* that produced the highest similarity updated the discrete template index to *k_t_* = *k_t_*_–1_ + *j*. The movement sequence phase *φ*(*t*) was then computed by normalizing *k_t_* by template length: *φ*(*t*) = *k_t_*/1000.

In the second step, muscle activity for channels artificially labeled as “missing” (to simulate paralysis) was reconstructed directly from the template at the newly decoded index *k_t_*. Although instantaneous phase was reported continuously, discrete indexing was necessary for the reconstruction process. Repeating these steps at each incoming data packet throughout the simulated data stream provided simultaneous estimates of instantaneous phase and reconstructed muscle activity.

### Differentiating between muscle activity templates

Dimensionality reduction analyses were performed on muscle activity templates to assess the feasibility of distinguishing among templates in real-time decoding scenarios. The analysis used the templates of the seven proximal muscles. When distal activity is reconstructed from proximal recordings, the proximal signals alone must identify the task, and the task-specific couplings among proximal muscles are visible and easy to separate, so a template can be selected or switched on the basis of those signals. Two methods were used: principal component analysis (PCA; linear method) and uniform manifold approximation and projection (UMAP; nonlinear method; parameters: 15 neighbors, minimum distance of 0.1, 2-dimensional embedding, random seed 12; see [36] for details). Following dimensionality reduction, two-dimensional histograms with 50 bins per axis were computed to quantify the overlap among template projections in linear and nonlinear spaces. Histograms were subsequently binarized into occupancy maps, wherein overlapping occupied bins indicated areas of intersection between templates. These intersections provided a quantitative measure of template separability, directly relevant to the feasibility of accurately selecting the appropriate task-specific template during real-time decoding.

### Statistical analyses

Normality of motor chunk time distributions was evaluated using the Anderson-Darling test. Differences between non-normal distributions were assessed using Kruskal-Wallis tests, followed by pairwise multiple comparisons with the Tukey-Kramer correction. Phase and muscle activity prediction accuracy was quantified using the root-mean-square error (RMSE), and the linear association between predicted and measured signals was assessed using Pearson’s correlation coefficient. Prior to PCA, the suitability of the data was assessed using Bartlett’s test of sphericity—to verify significant correlations among variables—and the Kaiser-Meyer-Olkin (KMO) measure—to ensure adequate sampling and variable interrelatedness. Statistical significance was set at α = 0.05 for all analyses. All procedures were executed using MATLAB R2024b (Statistics and Machine Learning Toolbox; MathWorks, Natick, MA, USA) and Python 3.13.5 (NumPy 2.5.4, SciPy 1.18.1, scikit-learn 1.9.1, umap-learn 0.5.12, factor_analyzer 0.5.1; the pinned environment is provided with the code).

## Results

A phase-based muscle activity decoder was constructed using population-averaged muscle-activation templates derived from EMG, kinematic, and FSR recordings obtained from three healthy participants (Subjects 1-3) who performed three standardized lateral object transfer tasks involving (1) gross finger flexion / extension (“grasp” task), (2) thumb opposition (“grip” task), and (3) wrist pronation / supination (“pronate” task) (see Materials and Methods). Testing was carried out on these initial participants, and validation was conducted with a fourth healthy participant (Subject 4), who had not contributed data for template generation, to determine whether phase predictions could be generalized beyond the original cohort. Subsequently, the predicted phase was employed to infer the activity of muscle groups (either of the proximal or distal arm), artificially labeled as missing to model paralysis. The simulated real-time performance of the decoder was then evaluated. The following sections detail these analyses and their outcomes.

Accelerometer (Fig 3a) and FSR (Fig 3b) signals were used to delineate the three principal motor chunks—Reach, Transfer, and Return—within each repetition. During the Reach chunk, the acceleration rose from baseline (‘movement starts’, Fig 3a), indicating that participants initiated forearm and wrist motion to engage the object. Concurrently, the FSR signal remained near zero until object contact, at which point the signal increased sharply, signifying object interaction (‘object grasped’, Fig 3b). The Transfer chunk began once the object was grasped and continued as long as the subject maintained contact with the FSR. A sharp drop in the FSR signal denoted object release (‘object released’, Fig 3b), marking the end of Transfer and the onset of Return chunk. The subsequent decrease of the accelerometer to baseline indicated cessation of the movement sequence (‘movement ends’, Fig 3a). The observed narrow standard-deviation bands (Fig 3a-b) reflected skilled upper-limb control and consistent task execution.

**Fig 3.**
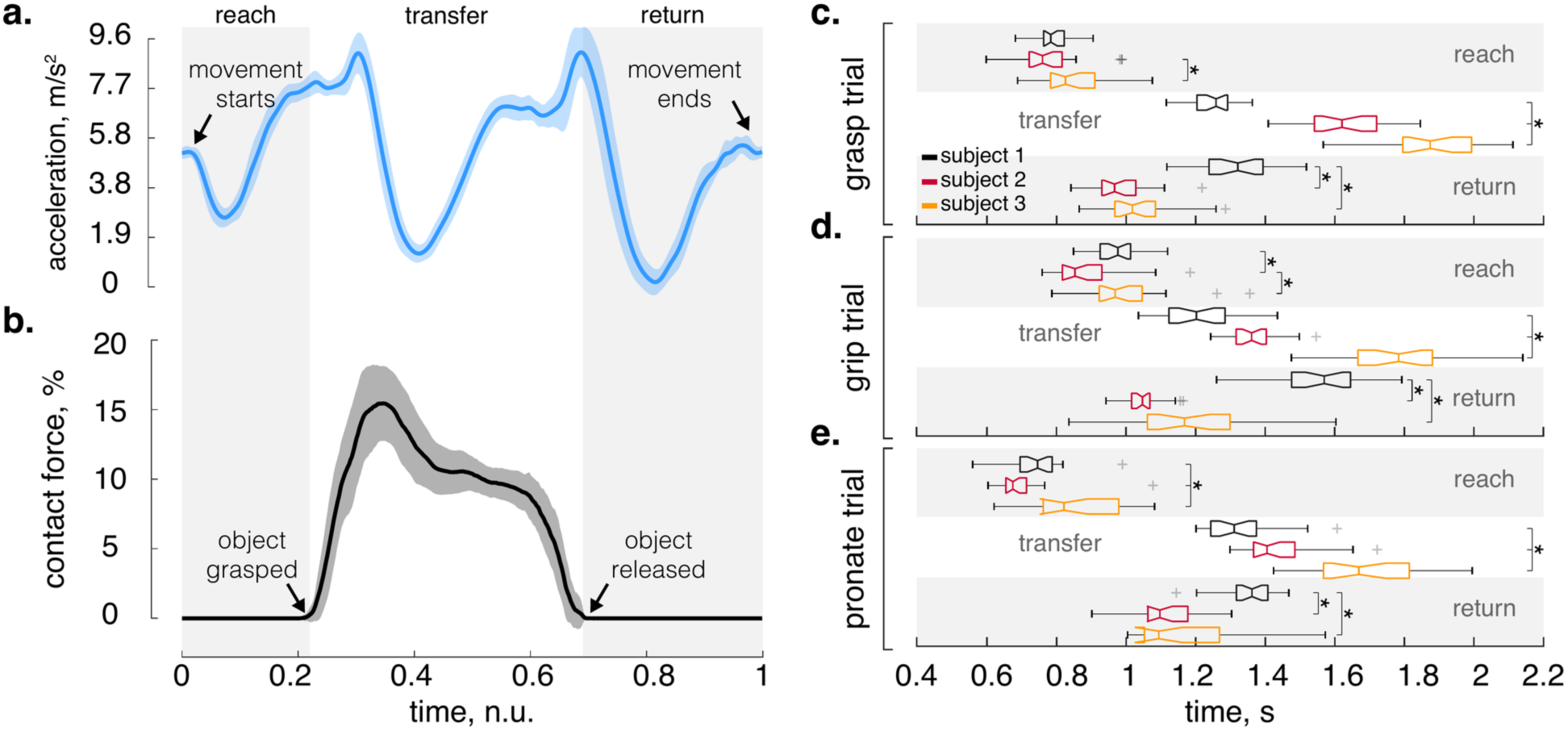
Sensor-based event identification and temporal variability of motor chunks. (**a**) Mean acceleration along one axis of the accelerometer (solid line) ± s.d. (shaded area) across 25 repetitions for a representative subject, recorded from a sensor placed over the FDS muscle. Deviation from the stable baseline acceleration indicates movement sequence initiation (‘movement starts’), while the return to baseline signifies movement sequence cessation (‘movement ends’). The alternation of acceleration and deceleration corresponds to three motor chunks—Reach, Transfer, and Return. (**b**) Corresponding mean FSR signal (solid line) ± s.d. (shaded area), expressed as a percentage of the sensor’s full-scale capacity (% FS), from the manipulated object for the same subject and repetitions. The rapid increase in force indicates engagement with the object (‘object grasped’), and the sharp decrease indicates cessation of object contact (‘object released’). (**c-e**) Boxplots illustrating the distribution of absolute durations for the Reach, Transfer, and Return motor chunks for each of the three subjects and tasks used in template generation. Despite pacing cues from a metronome, considerable inter-subject variability in chunk durations was observed, reflecting inherent biological variation and individual differences in synchronizing movements precisely with an external beat. Boxplots display the median (notch), interquartile range (IQR, box edges), with whiskers extending to 1.5 times the IQR from the box, and outliers plotted individually (crosses). Asterisks mark pairwise differences in chunk duration between subjects (Kruskal-Wallis test followed by Tukey-Kramer post hoc comparisons, p < 0.01).

Despite metronome pacing, there was notable inter-subject variability in the durations of Reach, Transfer, and Return (Fig 3c-e) chunks. Such variability arose from individual differences in preferred movement speeds and an inability to precisely follow external pacing (see [37] for a similar observation in an able-bodied cohort). To mitigate this variability across participants, data from each repetition were rescaled (see Materials and Methods) so that the Reach, Transfer, and Return chunks occupied predetermined, average proportions of the total movement sequence—namely, 22.99%, 45.06%, and 31.95% for the grasp task; 25.92%, 39.78%, and 34.30% for the grip task; and 22.08%, 42.91%, and 35.01% for the pronate task, respectively. This normalization aligned the motor chunks across subjects, permitting consistent analysis of the temporal evolution of muscle activity and creation of muscle activity templates.

Examination of muscle activity templates demonstrated that certain muscles exhibited consistent activation patterns across tasks, whereas others displayed clear task-specific modulation (Fig 4). Anterior deltoid (DeltA) activation remained relatively uniform across tasks. In contrast, middle deltoid (DeltM) activity increased distinctly at the transition from Transfer to Return during the pronate task. Posterior deltoid (DeltP) activation differed as early as the Reach chunk; there was elevated activity during the grasp task and comparatively lower DeltP activation during the pronate task. Similar to DeltM, DeltP activity also increased during the Transfer-to-Return transition in the pronate task.

**Fig 4.**
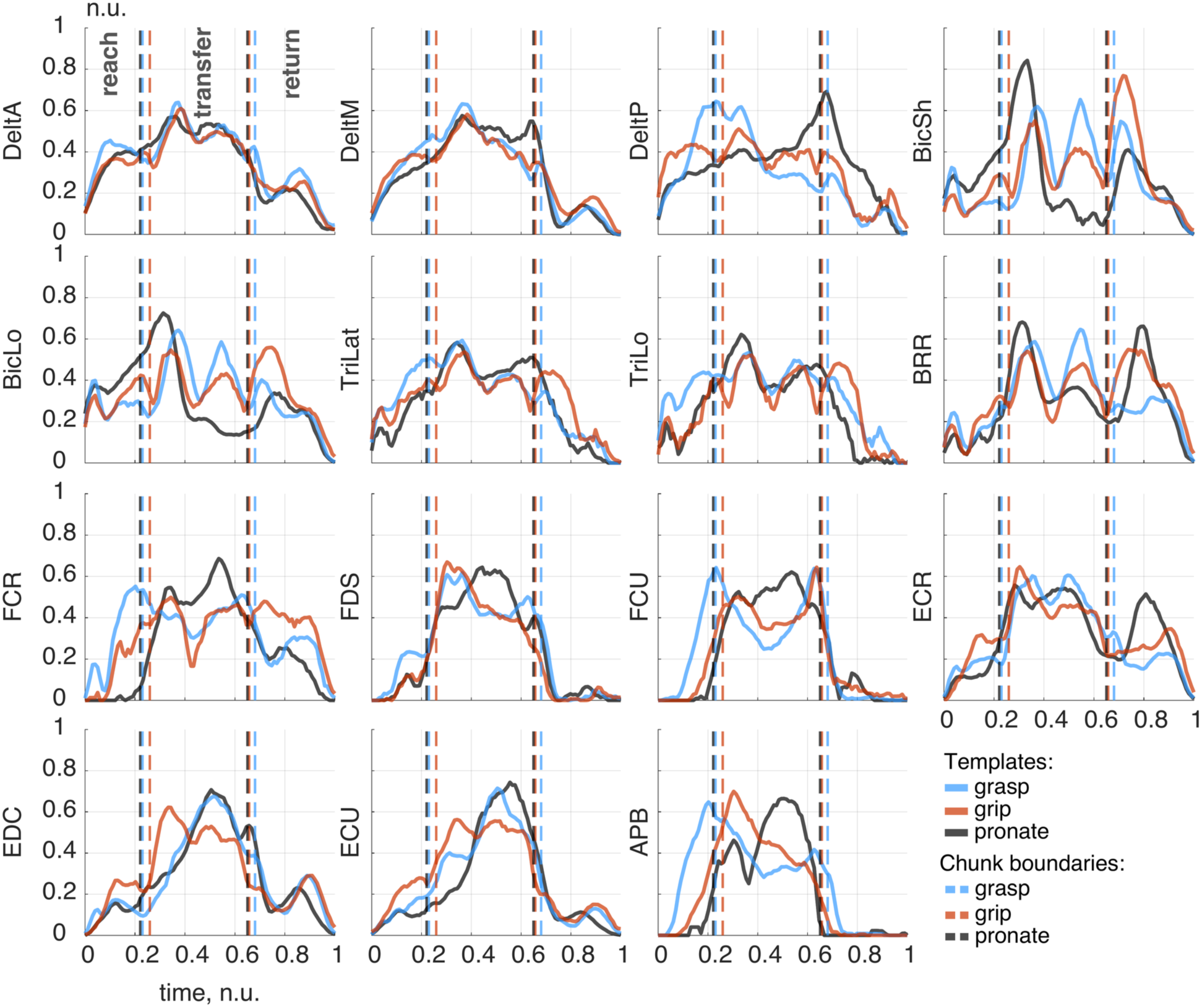
Average muscle-activity templates for 15 upper-limb muscles recorded as participants performed three lateral object transfer tasks—”grasp” (blue trace), “grip” (red trace), and “pronate” (black trace). *The vertical axis shows muscle activation in normalized units (n.u.) from 0-1*, and the horizontal axis shows normalized time from movement-sequence onset (0) to its completion (1). Each plot corresponds to a distinct muscle: DeltA (anterior deltoid), DeltM (middle deltoid), DeltP (posterior deltoid), BicSh (biceps short head), BicLo (biceps long head), TriLat (triceps lateral head), TriLo (triceps long head), BRR (brachioradialis), FCR (flexor carpi radialis), FDS (flexor digitorum superficialis), FCU (flexor carpi ulnaris), ECR (extensor carpi radialis), EDC (extensor digitorum communis), ECU (extensor carpi ulnaris), and APB (abductor pollicis brevis). Vertical dashed lines in each color mark the average (normalized) Reach, Transfer, and Return chunk boundaries for a given task, after each repetition timing was rescaled to the same proportional durations. The grasp task required gross finger flexion-extension with a cube transfer, the grip task emphasized thumb opposition with a cylindrical weight, and the pronate task involved flipping (pronating) a plate over a barrier. Distinct amplitude and timing patterns highlight the varying mechanical demands imposed by the three tasks.

Biceps short (BicSh) and long heads (BicLo) demonstrated elevated activation during the Reach chunk in the pronate task. During Transfer, both heads exhibited a characteristic biphasic activation pattern. The amplitude of this pattern was scaled by object mass. The second activation peak was also notably lower during the pronate task compared with the other tasks. Additionally, there was higher biceps activation in the grip task during the Return chunk. Triceps (TriLat and TriLo) activation patterns were generally consistent across tasks, with two exceptions: (1) there was greater initial activation during the Reach chunk in the grasp task, and (2) there was prolonged activation during Return in the grip task. Brachioradialis (BRR) showed similar patterns to the biceps heads across tasks but exhibited distinctly lower activation during Return after participants carried the heavier cube in the grasp task.

Distal arm muscle activation templates highlighted differences between tasks more prominently. FCR and FCU demonstrated higher activation amplitudes during Reach in the grasp task. Conversely, there was minimal FCR activation in the pronate task. There were transient reductions in flexor activation observed during Transfer. In contrast, FDS increased its activation during Transfer. During the Transfer chunk of the pronate task, there was prominent activation of ECR, EDC, and ECU. The extensor peaks reappeared at the end of the Return chunk. Lastly, abductor pollicis brevis (APB) showed clear task-dependent and aperture-related activation differences during Reach. The most prominent difference was the biphasic modulation during the Transfer chunk of the pronate task.

Fig 5 illustrates the decoding of the normalized phase (scaled from 0 n.u. at movement sequence onset to 1 n.u. at movement sequence completion) using three distinct muscle-channel counts: all muscles (Fig 5, panels c-f, top row), proximal arm muscles only (7 muscles; panels g-j, middle row), and distal arm muscles only (8 muscles; panels k-n, bottom row). Data are presented for three “observed” subjects whose muscle activities informed the creation of decoding templates (Subjects 1-3), and for one “unobserved” subject (Subject 4) to validate generalizability. For clarity, Fig 5a schematically illustrates the phase progression of three consecutive movement sequences, each represented as a linear ramp (from 0 to 1 n.u.), with resting intervals shown as flat segments between sequences. Fig 5b shows how resting intervals were omitted from subsequent visualizations to isolate the temporal progression of movement phases and eliminate unrelated idle periods.

**Fig 5.**
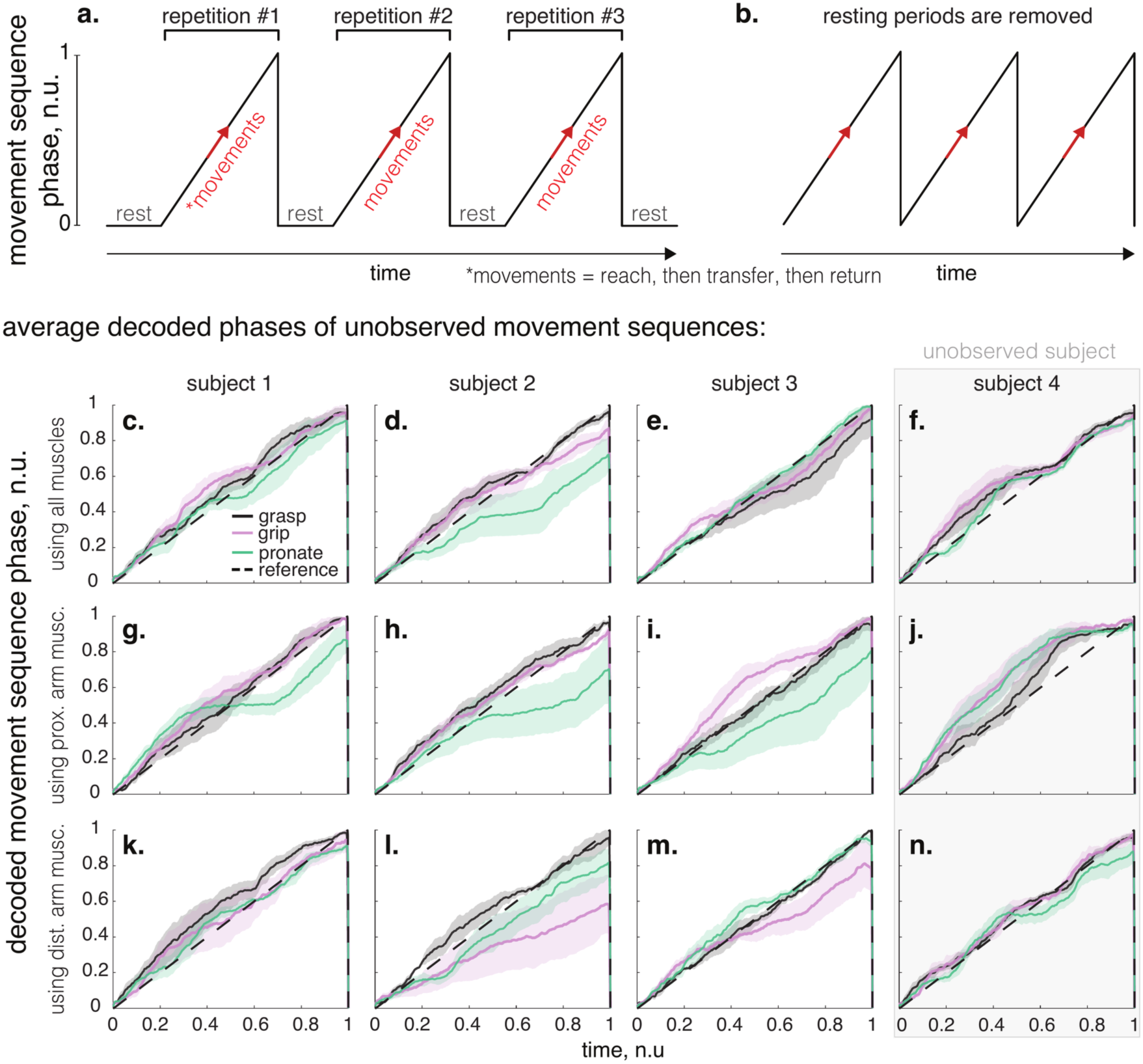
Inference of phase (0-1 n.u.) of the Reach-Transfer-Return movement sequence under different muscle-channel counts—using all available muscles (top row, panels **c-f**), only proximal arm muscles (middle row, panels **g-j**), or only distal arm muscles (bottom row, panels **k-n**). Panels **a** and **b** provide schematic representations of how resting segments are removed, leaving only the active interval where movement sequence progresses from phase 0 (onset) to phase 1 (completion). Colored traces in panels c-n display the averaged phase estimates across grasp (black), grip (pink), and pronate (green) tasks, plotted against a dashed diagonal indicating perfect phase prediction. The first three columns (c, g, k; d, h, l; e, i, m) show the phase decoder’s performance on *withheld* repetitions of the subjects whose data were included in constructing the EMG templates, while the final column (f, j, n) shows performance for an “unobserved” subject not involved in template generation. The close correspondence between the decoded trajectories and the diagonal demonstrates accurate phase tracking, and the similar performance in the unobserved subject underscores the generalizability of the population-averaged templates.

For Subjects 1-3, the decoding outcomes across tasks are displayed as average phase progressions—computed for 10 *withheld* repetitions—superimposed on a dashed diagonal representing perfect decoding accuracy (Fig 5c-e, g-i, k-m). Overall, when all muscle channels are considered, each subject’s decoded phases closely align with the diagonal (error distributions in Fig 6a). Although standard deviations were generally narrow, increases in some cases likely reflected subject-specific variations in muscle coordination across tasks. Importantly, the decoder’s accuracy extended to Subject 4; for this subject, the phase trajectories were computed and averaged for 25 repetitions (Fig 5f, j, n). Despite minor phase estimation deviations around mid-movement sequence—potentially arising from subtle individual differences in coordination or timing (for example, in some repetitions the hand returned behind rather than over the barrier, S1 Fig)—the decoded phase maintained proximity to the reference (see error distributions in Fig 6d).

**Fig 6.**
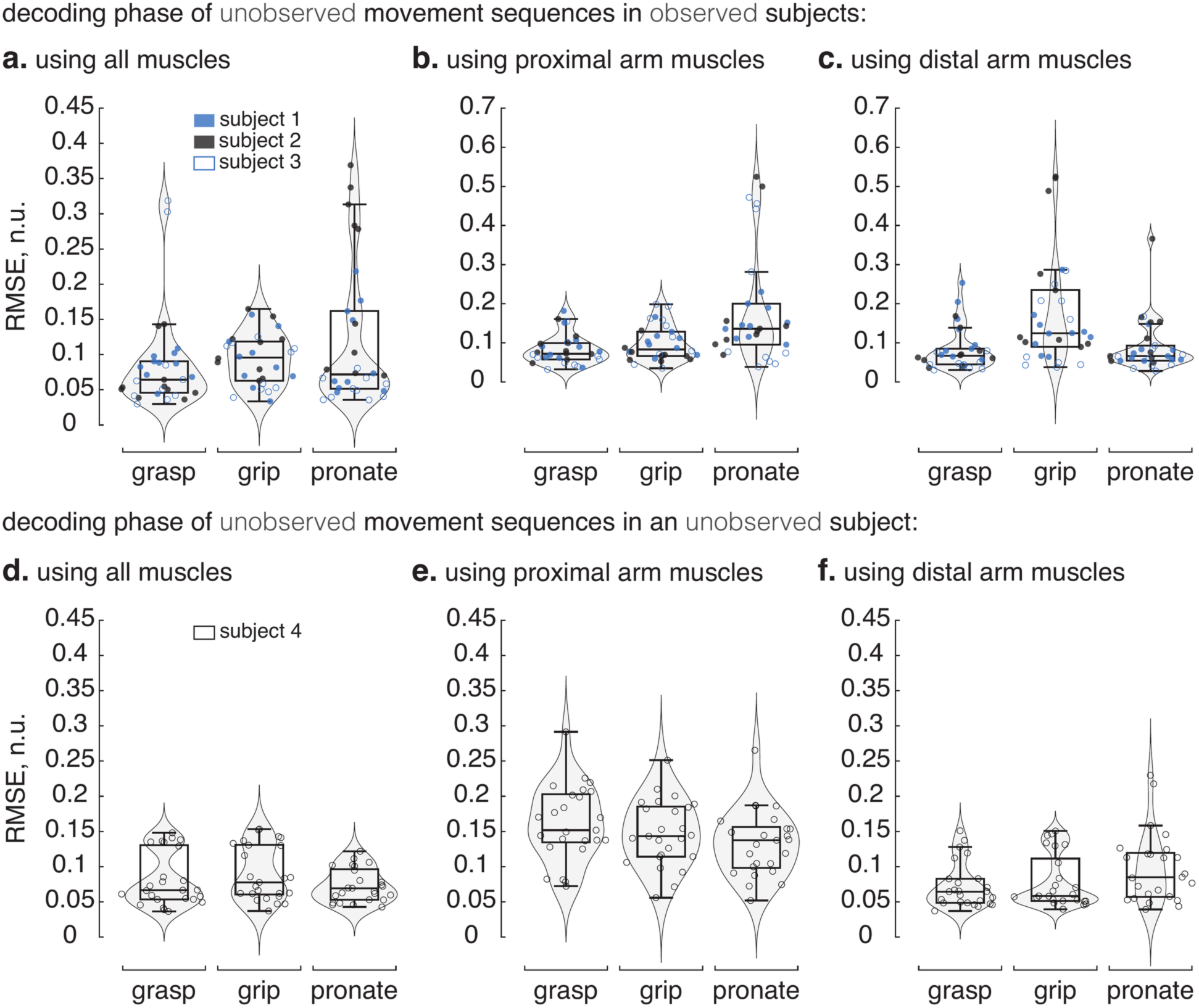
Phase-decoding errors (root-mean-square error, RMSE) for each of the three lateral object transfer tasks—grasp, grip, and pronate— shown as a function of channel count for observed and unobserved participants. Panels (**a-c**) present results for the three observed participants whose recordings contributed to building the population-averaged templates (performance is shown for 10 repetitions *withheld* from the templates), while panels (**d-f**) show performance for an unobserved participant (25 repetitions) who was not involved in template generation. In each row, decoding errors are shown for when (**a,d**) all available muscles, (**b,e**) only proximal muscles, and (**c,f**) only distal muscles were used. The violin plots visualize the probability distribution of errors, with overlaid boxplots denoting median, interquartile range, and outliers. Smaller RMSE values indicate closer alignment of the decoded movement phase with its true progression. Robust performance on the unobserved participant (bottom row) underscores the generalizability of the population-based templates.

Across tasks, decoding accuracy depended on channel count. For Subjects 1-3 (Fig 6a-c), using all muscle channels (Fig 6a) yielded relatively low RMSEs across grasp (0.085 ± 0.068 n.u.), grip (0.094 ± 0.036 n.u.), and pronate tasks (0.121 ± 0.100 n.u.). Restricting decoding to proximal muscle channels (Fig 6b) provided comparable accuracy for grasp (0.083 ± 0.038 n.u.) and grip tasks (0.098 ± 0.044 n.u.), but increased errors for the pronate task (0.184 ± 0.145 n.u.), reflecting the importance of forearm musculature in rotational movements of the distal arm. Conversely, decoding using distal channels alone (Fig 6c) maintained accuracy for grasp (0.082 ± 0.054 n.u.) and pronate tasks (0.089 ± 0.065 n.u.), but increased errors during the grip task (0.174 ± 0.136 n.u.), indicating that grip-task decoding substantially benefits from proximal muscle signals.

In the unobserved participant (Fig 6d-f), the dependence on channel configuration differed from that in Subjects 1-3. Although using all channels (Fig 6d) resulted in consistently low RMSEs (grasp: 0.081 ± 0.038 n.u., grip: 0.090 ± 0.039 n.u., pronate: 0.073 ± 0.023 n.u.), decoding using distal-only channels (Fig 6f) slightly outperformed the all-channel condition in both grasp (0.072 ± 0.032 n.u.) and grip tasks (0.079 ± 0.038 n.u.). However, distal-only decoding was less accurate for the pronate task (0.098 ± 0.050 n.u.), indicating task-specific reliance on both proximal and distal musculature. In contrast, proximal-only recordings (Fig 6e) consistently yielded higher errors (grasp: 0.161 ± 0.053 n.u., grip: 0.148 ± 0.047 n.u., pronate: 0.133 ± 0.047 n.u.). Overall, each task displayed a distinct dependence on channel configuration, yet accurate decoding remained achievable across all tested channel counts.

The predicted movement phase was used to index the task-specific templates to estimate the activity of muscles labeled as “missing” to emulate the effects of paralysis. Predictions for distal muscle activity (which relied on proximal muscle channels) were evaluated first. Across the three tasks, the predicted muscle activity traces showed notable variation in quality, as evidenced by the correlation coefficients (r) and RMSEs (Fig 7). Predicted activity of brachioradialis (BRR), which participates in elbow flexion and influences forearm rotation, exhibited moderate agreement with empirical signals during grasp (r=0.51, RMSE=0.18) and grip (r=0.53, RMSE=0.14) tasks, and more pronounced alignment in the pronate task (r=0.72, RMSE=0.13). If these decoded BRR patterns were employed to deliver electrical stimulation, one would expect reasonably synchronized assistance for forearm pronation, although the moderate correlations in grasp and grip might still necessitate calibration to refine elbow flexion support during those tasks.

**Fig 7.**
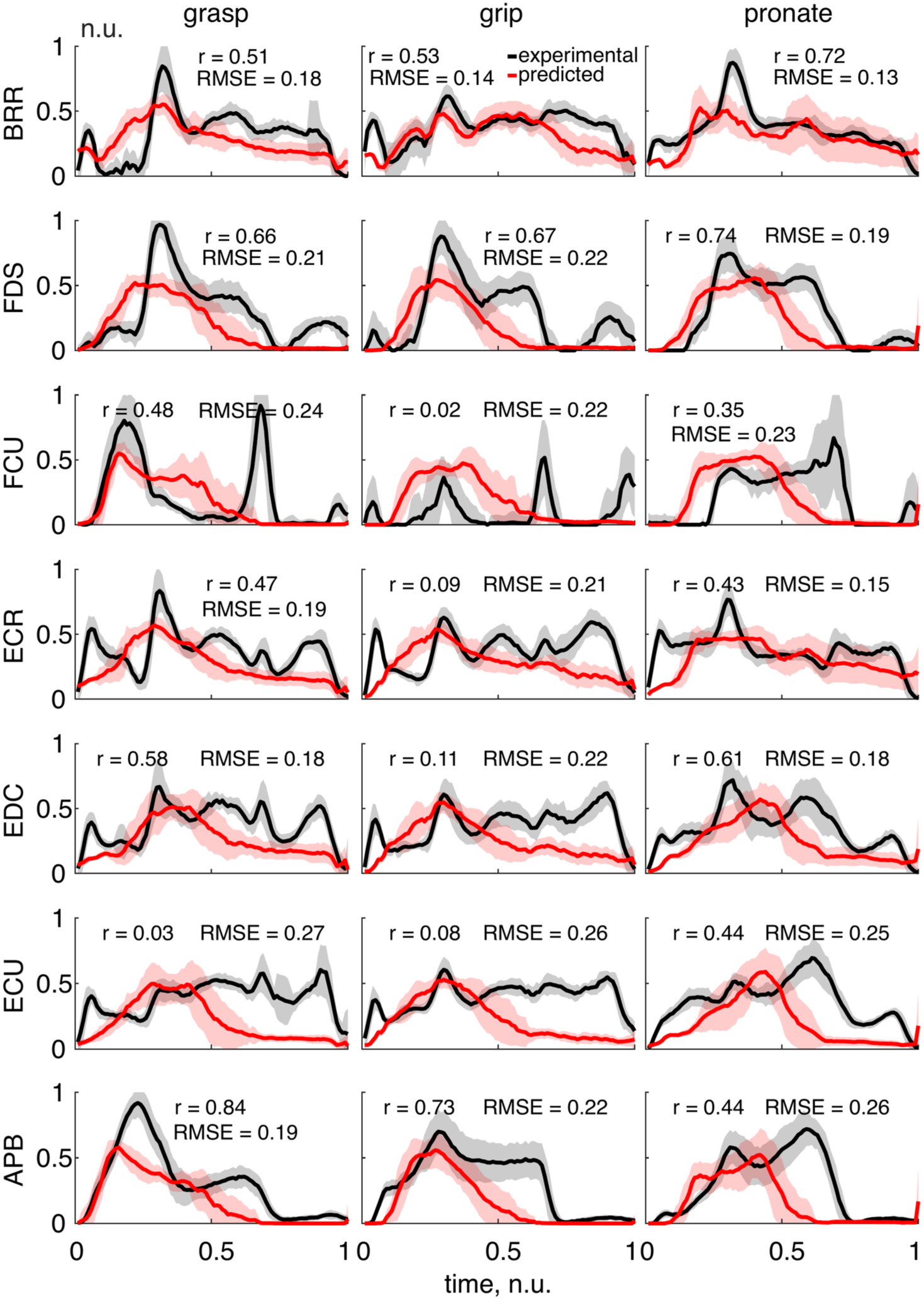
Predicted versus experimentally recorded *distal* muscle activity in the unobserved participant (Subject 4) during three lateral object transfer tasks—grasp, grip, and pronate—for brachioradialis (BRR), flexor digitorum superficialis (FDS), flexor carpi ulnaris (FCU), extensor carpi radialis (ECR), extensor digitorum communis (EDC), extensor carpi ulnaris (ECU), and abductor pollicis brevis (APB) (FCR was excluded; see Data exclusion). In each subplot, the black trace and accompanying shaded region represent the mean and standard deviation of the experimentally measured EMG activity, while the red trace and shaded region depict the decoded (predicted) output. The correlation coefficient (r) and root mean square error (RMSE) values, computed between the mean measured and mean predicted traces, are displayed within each subplot. Higher correlation and lower RMSE generally indicate stronger agreement between decoded and actual muscle activity time courses. The time axis (0-1) corresponds to a normalized movement duration for each task, and the vertical is normalized muscle activity (n.u.).

Predicting the activity of muscles stabilizing the wrist emerged as a major challenge, particularly for FCU and ECR, whose predicted signals deviated substantially in specific tasks. FCU showed a moderate correlation (r=0.48) only in grasp, with poor alignment for the grip (r=0.02) and pronate (r=0.35) tasks, suggesting that the timing and amplitude demands of wrist flexion in grip were missed by the decoder. ECR performed similarly poorly in the grip task (r=0.09), while providing moderate congruence during the grasp (r=0.47) and pronate (r=0.43) tasks. These observations imply that any FES regimen relying on these decoder outputs might misapply wrist flexion or extension forces when grasping or gripping, potentially compromising object handling and overall hand-wrist coordination. ECU also yielded low correlations (≤0.08) in both grasp and grip, though its moderate pronate correlation (r=0.44) suggests a partial salvage of predictive fidelity for tasks demanding forearm rotation.

Regarding muscles controlling finger movements, FDS predictions exhibited robust performance (r≥0.66) across the grasp, grip, and pronate tasks, pointing to reliable detection of the underlying flexion demands required for digital closure around various objects. Although the RMSE values (≈0.20) indicate amplitude offsets, the overall alignment in shape and timing suggests that using these decoded signals to stimulate finger flexors could yield suitably synchronized closure forces. Meanwhile, EDC predictions were reliable for pronate (r=0.61) and to a lesser extent for grasp (r=0.58) tasks but failed to track the extension demands during grip (r=0.11). One might infer that finger extension is not modeled effectively in tasks involving strong finger flexion until the object is released.

APB, responsible for thumb opposition and stabilization, showed high decoding fidelity for grasp (r=0.84) and grip (r=0.73). Accurate tracing of its activation patterns in these two tasks would likely translate into precise electrical stimulation for stabilizing the thumb during object pick-up, transfer, and placement. Accuracy of APB activity prediction declined in the pronate task (r=0.44), which is not surprising given that thumb abduction is less heavily emphasized when the primary objective is flipping the plate through forearm rotation.

Overall, these results delineate a pronounced discrepancy between muscles whose decoded activity closely mirrors their actual physiological signals—namely FDS and APB—and those for which the decoder appears less accurate at capturing the nuances of activation, such as FCU, ECR, and ECU. While FDS and APB-based stimulation would likely produce more consistent and functionally meaningful hand movements, the comparatively weak predictions for wrist stabilizers underscore the need for refined decoding or complementary strategies to reliably support wrist positioning under an FES.

Predictions for proximal muscle activity (relying on distal arm muscle channels) were evaluated next (Fig 8). Recorded and predicted DeltA activity matched in all three tasks, as indicated by correlations of 0.88 or higher and RMSE values in the 0.08-0.11 range. This consistency suggests that the decoder captures the timing and amplitude necessary for forward shoulder positioning during reach and transfer. If these signals were used for FES, the user would likely receive well-synchronized anterior deltoid support, aiding reliable arm elevation and stabilization. DeltM— recorded poorly in Subject 4—showed moderate performance in the grasp task (r=0.64, RMSE=0.16) but fell to negative correlations for grip and pronate (r=−0.18 for both), accompanied by comparatively high RMSE (up to 0.34). Posterior deltoid (DeltP) achieved near-perfect agreement in grasp (r=0.97, RMSE=0.08) and maintained strong correspondence for the grip task (r=0.85, RMSE=0.13), yet dropped to moderate accuracy in the pronate task (r=0.48, RMSE=0.25). This decline suggests that, while the decoder effectively recognizes posterior deltoid activation during forward lifting and holding tasks, it struggles to track the more subtle changes in scapular retraction or shoulder rotation associated with plate-rotating motions. Under FES, these weaknesses may manifest as poorly timed or insufficient posterior shoulder engagement during dynamic pronation movements.

**Fig 8.**
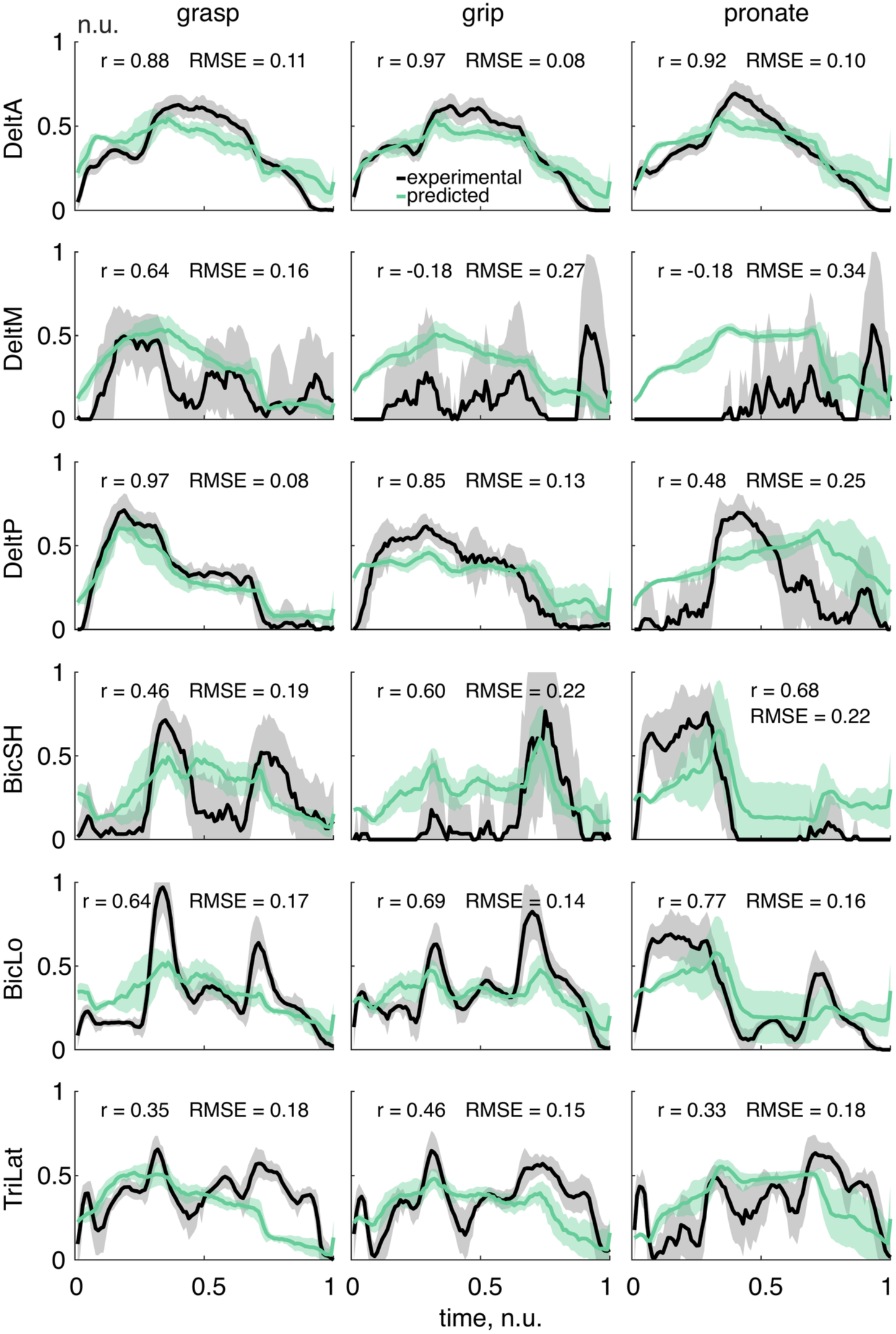
Predicted versus experimentally recorded *proximal* muscle activity in the unobserved participant (Subject 4) during three lateral object transfer tasks—grasp, grip, and pronate—for anterior deltoid (DeltA), middle deltoid (DeltM), posterior deltoid (DeltP), biceps short head (BicSh), biceps long head (BicLo), and triceps lateral head (TriLat) (TriLo was excluded; see Data exclusion). In each subplot, the black trace and accompanying shaded region represent the mean and standard deviation of the experimentally measured EMG activity, while the green trace and shaded region depict the decoded (predicted) output. Other descriptions are similar to those in Fig 7.

BicSh and BicLo both exhibited moderate-to-strong correlations, although BicLo activity predictions consistently outperformed BicSh across tasks, partly due to greater signal quality (visual inspection). BicLo’s correlation values ranged from r=0.64 (grasp task) to r=0.77 (pronate task), implying a fairly accurate capture of elbow flexion activity. BicSh, while still moderately accurate (r=0.46-0.68), showed broader amplitude mismatches, especially in grip and pronate tasks (RMSE=0.22). In an FES setting, BicLo’s higher prediction fidelity would translate to more precise support of elbow flexion, whereas BicSh’s less accurate signals might generate mismatched levels of force, affecting the consistency of elbow control.

TriLat was among the least accurately predicted of these muscles, with correlation coefficients of 0.35 in grasp, 0.46 in grip, and 0.33 in pronate tasks. Such modest alignment indicates the decoder’s limited effectiveness at detecting the onset and magnitude of elbow extension. Any stimulation strategy reliant on this predictive signal would risk erratic or mistimed activation, which could undermine the user’s ability to smoothly extend or stabilize the elbow against gravity during object manipulation.

In sum, DeltA, DeltP, and BicLo predictions matched recorded muscle activity most coherently, suggesting these signals could be harnessed with relative confidence to drive stimulation-based support for shoulder and elbow motions. In contrast, DeltM, BicSh, and TriLat predictions showed greater variability, phase mismatches, or limited amplitude accuracy, indicating that further decoding refinements are needed to integrate these muscles effectively into an FES paradigm.

Beyond decoding accuracy, a key consideration for any decoder-stimulator paradigm is its suitability for real-time applications. To assess whether our decoding algorithm can operate under real-time constraints, we measured the execution time of each stage. The processing steps (see Materials and Methods) on average required 0.8 ± 0.3 ms (mean ± s.d.; Fig 9a) per 14 ms packet. Decoding the movement sequence phase introduced similarly minimal delays, averaging 1.3 ± 1.1 ms per 14 ms packet when all muscle channels were used, 0.9 ± 1.1 ms with proximal arm muscles only, and 1.0 ± 1.2 ms with distal arm muscles only (Fig 9b). Collectively, these processing and decoding latencies remained within low-millisecond ranges (<13.5 ms packet duration of the Delsys Trigno® Wireless Biofeedback System), indicating that our decoding method imposes negligible computational overhead and is therefore well-suited for applications requiring real-time performance. (Notably, infrequent high-latency outliers arose from occasional TCP/IP packet-transfer delays between the acquisition and decoding computers / systems, not from the decoding algorithm).

**Fig 9.**
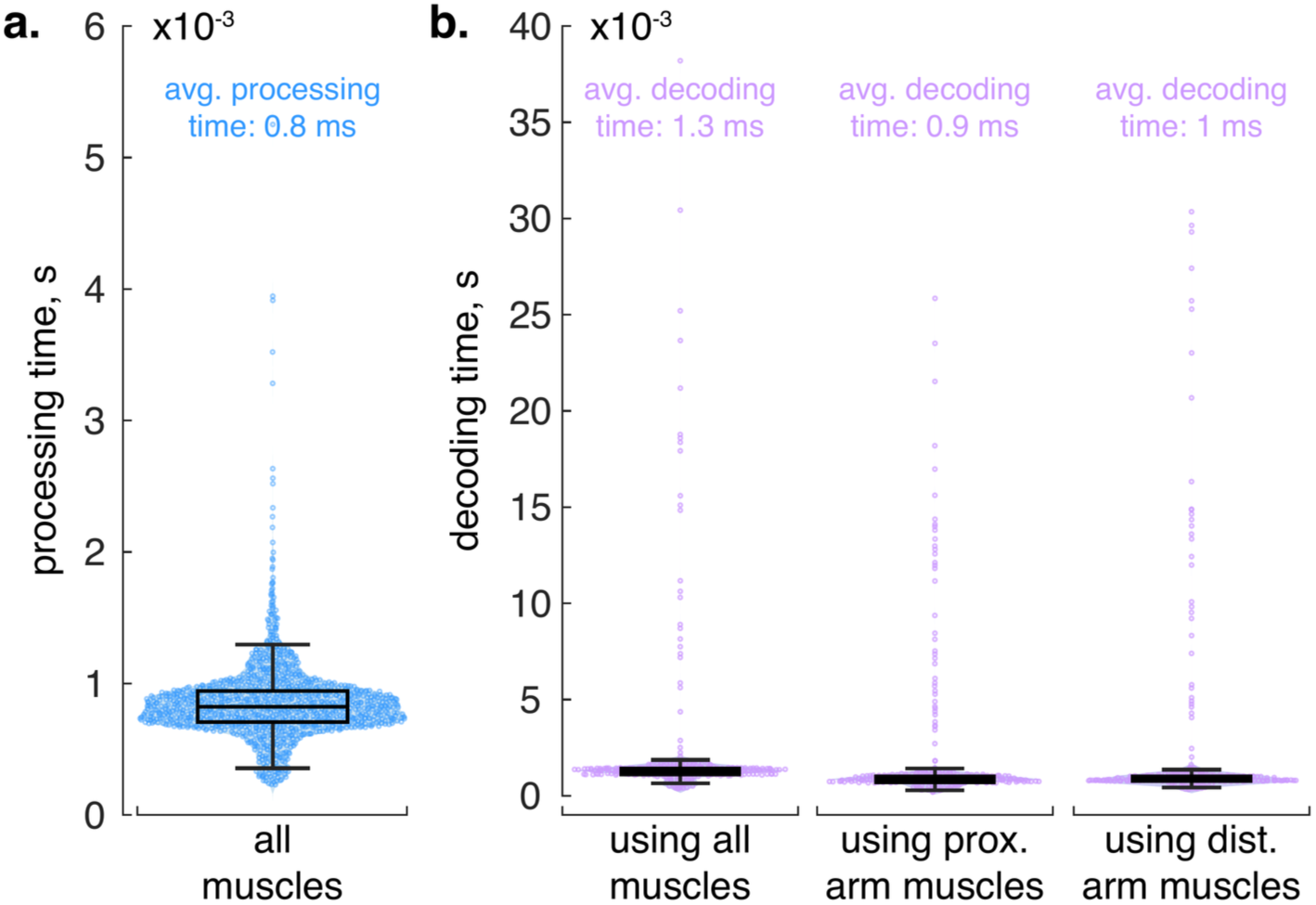
Processing and decoding times. (**a**) Box and swarm plots show the time required to execute the EMG processing pipeline for a 14 ms data packet containing muscle activity recordings from all channels. The boxplot denotes median (horizontal line), interquartile range (box edges), and outliers; the mean processing time is approximately 0.8 ms. (**b**) Average decoding delays for three different muscle channel selections— using all muscles (1.3 ms average), only proximal muscles (0.9 ms average), or only distal muscles (1.0 ms average). Each boxplot depicts the distribution of decoding latencies, with swarm plots demonstrating data points; all distributions (a-b) contain 5,000 data points. Lower values indicate faster updates in both processing and decoding, supporting real-time estimation of the movement-sequence phase. *The few high-latency outliers arose from intermittent TCP-IP packet-transmission delays between acquisition and decoding computers / systems, not from the decoding algorithm*.

Another important consideration is the ability to distinguish among multiple task-specific muscle-activation templates in real time. Fig 10 shows the mean activation templates of the seven proximal muscles for each task projected into both PCA (Bartlett’s test, *χ*^2^ = 31602.85, p<0.01; KMO=0.62-0.81) and UMAP spaces to illustrate the feasibility of template discrimination. In the PCA projection (Fig 10a), grasp and grip task-related projections shared 46 bins (more explicitly, intersection points; see Materials and Methods), grasp and pronate tasks shared 48, and grip and pronate tasks shared 28, indicating that a linear approach does not clearly separate the task-related templates. By contrast, the UMAP embedding (Fig 10b) produced no shared bins between grasp and either grip or pronate tasks, and only two bins between grip and pronate tasks. This near-complete separation suggests that a nonlinear method (UMAP or other) can provide greater specificity for identifying the correct template in real time. In practice, ongoing EMG signals could be compared to each stored template (e.g., via dot products) to determine which task is being performed, enabling a switch from one decoding mode to another when muscle-activation patterns align most strongly with that template. Although a switching protocol was not implemented in this study, the minimal overlap in the UMAP domain supports the viability of such an approach.

**Fig 10.**
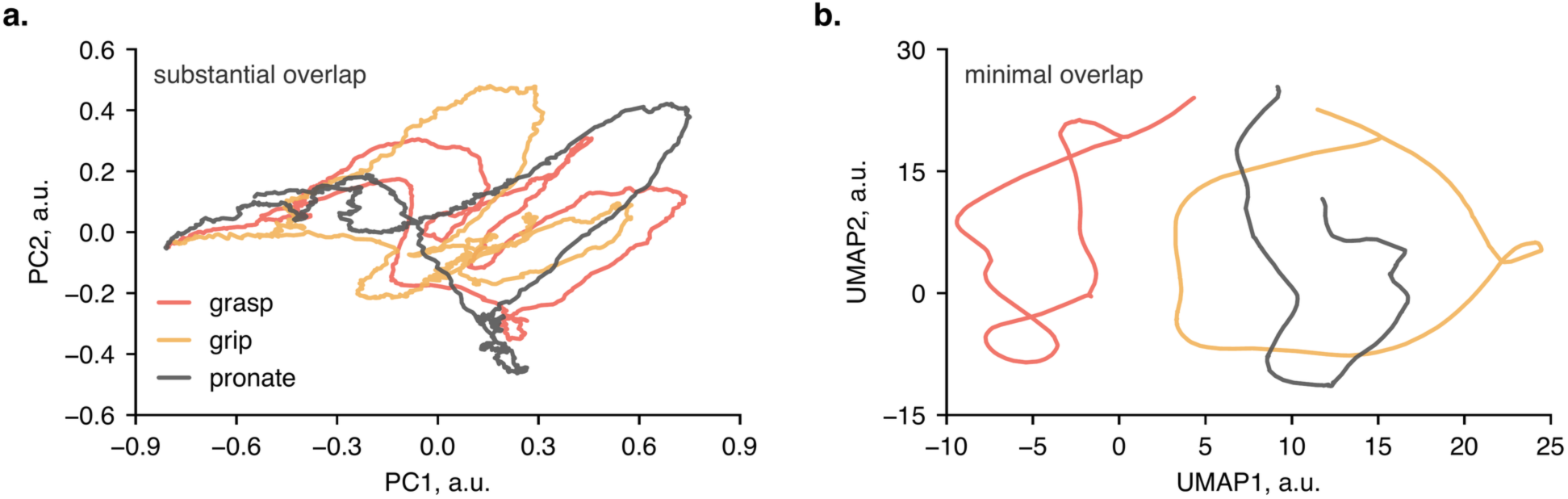
Visualization of the mean activation templates of the seven proximal muscles for grasp (red), grip (orange), and pronate (black) tasks projected into two distinct low-dimensional spaces. (**a**) Principal component analysis (PCA) reveals considerable overlap among the three trial types, consistent with a histogram-based analysis showing 46 shared bins (points of intersection; see Materials and Methods) between grasp and grip tasks, 48 between grasp and pronate tasks, and 28 between grip and pronate tasks. This indicates that a linear transformation alone does not reliably separate each task’s muscle-activation pattern. (**b**) Uniform manifold approximation and projection (UMAP) produces well-separated arrangements of the same data, with zero bins overlapped for grasp and grip or grasp and pronate tasks, and only two overlapped bins between grip and pronate tasks. Such a nonlinear mapping suggests the feasibility of a real-time template switching strategy, in which projections of ongoing muscle activation signals are compared with projections of templates (e.g., via dot products) to determine which trial type is being executed. Axes are in arbitrary units (a.u.).

## Discussion

In this study, we demonstrate that missing muscle activity underlying different functional tasks can be reconstructed under simulated real-time constraints using a sparse set of EMG signals and a small library of muscle-activity templates indexed by a single continuously updated phase variable. Using EMG from 15 upper-limb muscles, we created task-specific templates from three able-bodied participants and tested them in a fourth participant under two simulated denervation scenarios: (i) retaining only distal arm EMG channels while hiding proximal channels, and (ii) the reverse. With just seven to eight remaining channels, the decoder reliably tracked movement phase (median RMSE < 0.17 n.u.; 0 = task start, 1 = task end), accurately reconstructed withheld muscle activity in several key muscles (RMSE 0.08-0.34, correlations up to 0.97) and operated with ∼2 ms delays. Nonlinear dimensionality reduction indicated clear separation among templates for the three object-transfer tasks, suggesting feasibility for automatic task recognition. These results demonstrate that our approach can rapidly and with “good-enough” accuracy infer missing muscle activity, providing a potential practical solution for real-time FES or robotic orthosis systems aimed at restoring arm function.

Prior to this work, multiple studies showed that a related strategy, relying on low-dimensional muscle activation patterns—termed muscle synergies—can recover missing muscle activation patterns with high fidelity. For example, extracting subject-specific synergies from recorded muscles enabled accurate prediction of the EMG patterns for novel, untrained hand postures [38]. Across diverse tasks and populations (able-bodied, stroke), as few as ∼4 synergies often suffice to reconstruct full-arm muscle activation patterns, accounting for >90% of the variance [39,40]. Building on this concept, recent methods combine synergy extrapolation with musculoskeletal modeling to estimate unmeasured muscle signals. In a healthy-arm study, a synergy-driven EMG model predicted deep shoulder or elbow muscle activations with up to R≈0.99 agreement when only superficial muscles were measured [18]. Even with multiple simultaneous missing muscles, the approach retained high accuracy (often >90% correlation) in estimating the absent activations [19]. Because some of these algorithms have yet to achieve real-time performance or have not been proven to scale when signals from several synergistic muscles are concurrently unavailable—as in cases of severe paralysis—their adoption in practical assistive-device control remains limited.

Alternate strategies have leveraged physiological modeling and optimization to predict missing EMG data. A hybrid neuromusculoskeletal model that personalized unmeasured inputs via static optimization was shown to substantially improve joint torque tracking over standard EMG-driven simulations [41]. Similarly, an EMG-driven forward dynamics approach using a neural co-contraction model reduced muscle force estimation errors by ∼25% when unmeasured muscles were accounted for [42]. Notably, a direct comparison found that synergy-extrapolation outperforms classical static optimization for concurrent prediction of multiple missing muscle signals (halving amplitude errors and quadrupling correlation coefficients) [43]. However, purely data-driven synergy fits can degrade in accuracy without some knowledge of the missing inputs— initial studies required adjusting synergy weights using the very signals meant to be predicted. To address such limitations, emerging techniques typically resort to unintuitive (yet effective) deep learning-based solutions (e.g., see [44] for convolutional neural network-based decoder achieving ∼98% of signal variance explained and an RMSE of about 0.11 in EMG reconstructions).

In parallel, brain-computer interface prostheses now bypass the need to rely on residual muscle activity entirely: in one study, a brain-controlled FES system activated 36 paralyzed arm muscles, enabling a tetraplegic subject to drink from a cup in 11/12 trials by decoding and reconstituting his intended muscle activity [45]. In the same vein, even individuals born without a limb can volitionally produce distinct muscle activation patterns for the missing limb—children with congenital below-elbow deficiency consistently produced at least five distinct residual-muscle activation patterns, imaged with ultrasound, corresponding to hand movements they attempted [46].

Unlike synergy-based or statistical decoders—which express muscle activity as a weighted sum of static patterns and must continually solve for those weights—our approach models each movement as progression along a pre-established template parameterized by a single phase variable. Real-time operation is reduced to two steps: (1) map the incoming sparse EMG onto a task-specific template to obtain the current phase in the movement sequence, and (2) read out the complete muscle activations corresponding to that phase. Because no iterative optimization is required, the method produces millisecond-latency reconstructions that remain accurate for several functionally important muscles, making it well suited for control of assistive technologies.

Our study deliberately adopted a tightly controlled, proof-of-concept design to establish technical feasibility before pursuing broad clinical translation, and that choice introduces several methodological constraints. First, muscle-activity templates were derived from neurologically intact adults, with denervation emulated by masking selected EMG channels. This approach decouples decoder performance from the highly variable signal characteristics seen after injury, yet it leaves open the question of how well the method will transfer to individuals with actual neuromuscular impairment (in those cases, templates could be generated from each participant’s contralateral, unaffected limb). Second, surface electrodes were applied in a single, fixed placement per participant to maximize within-session consistency; in real-world practice, however, electrode positions often shift across sessions or must be customized to anatomy and residual function. Third, although our dimensionality-reduction analysis indicates that automatic template switching is feasible, we evaluated decoding accuracy only within isolated, single-task trials; confirming robustness during rapid task-to-task transitions remains an essential next step. Variants of the same task, such as returning the hand behind rather than over the barrier (S1 Fig), change the underlying muscle activity and lowered decoding accuracy in the affected repetitions, so a template library would need to cover such variants.

Another limitation of the study is that the muscle-activity templates were derived from only three human participants. Nevertheless, converging evidence indicates that a small-*N*, many-repetitions design can yield reliable representations of motor behavior. Across both human and animal EMG studies, low-dimensional muscle synergies have been recovered from cohorts as small as three to ten individuals, provided that within-subject sampling is dense [25,26,39,47]. Likewise, population-level neural manifolds measured with intracortical electrodes are typically characterized in two or three macaques yet remain remarkably stable across animals and sessions [48,49]. As Smith and Little [50] argue, statistical power accrues more from numerous repeated measures than from the sheer number of unique individuals—particularly when the signal of interest (here, the participant’s phase-indexed muscle-activations) is highly stereotyped and therefore exhibits a high within-subject signal-to-noise ratio. Our design follows this line of argumentation since each participant performed 25 repetitions of each object-transfer task, allowing the phase-template constructing algorithm to average out repetition-level noise. Holding out a fourth participant for validation further showed that templates built from three subjects generalized to an unseen individual, implying that the extracted templates capture task-relevant structure shared across individuals rather than subject-specific variability. Finally, the field routinely accepts “two-to-three-monkey” datasets as sufficient for establishing core principles of motor-cortex dynamics because cross-subject agreement—not sample size per se—is the critical criterion (e.g., see [51]). On that basis, our human *N* represents a comparable methodological choice, although larger cohorts will be necessary to probe factors such as limb dominance and coordination impairment.

## Acknowledgments

We thank Drs. Valeriya Gritsenko, Anna Korol, Shadman Tahmid, and others for their feedback.

## Supporting information

**S1 Fig.**
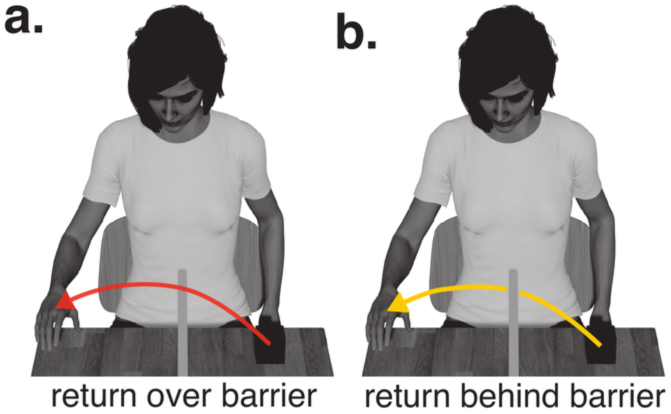
Two strategies for the Return chunk. (a) Return of the hand over the barrier, the instructed movement. (b) Return behind the barrier, which participants adopted in some repetitions. The muscle activity underlying the two movements differs, which lowers the decoding accuracy of the affected repetitions and contributes to the deviations of the decoded phase around the middle of the movement sequence.

